## Supporting information for "Neuropeptide Bursicon and its receptor mediated the transition from summer-form to winter-form of *Cacopsylla chinensis*"

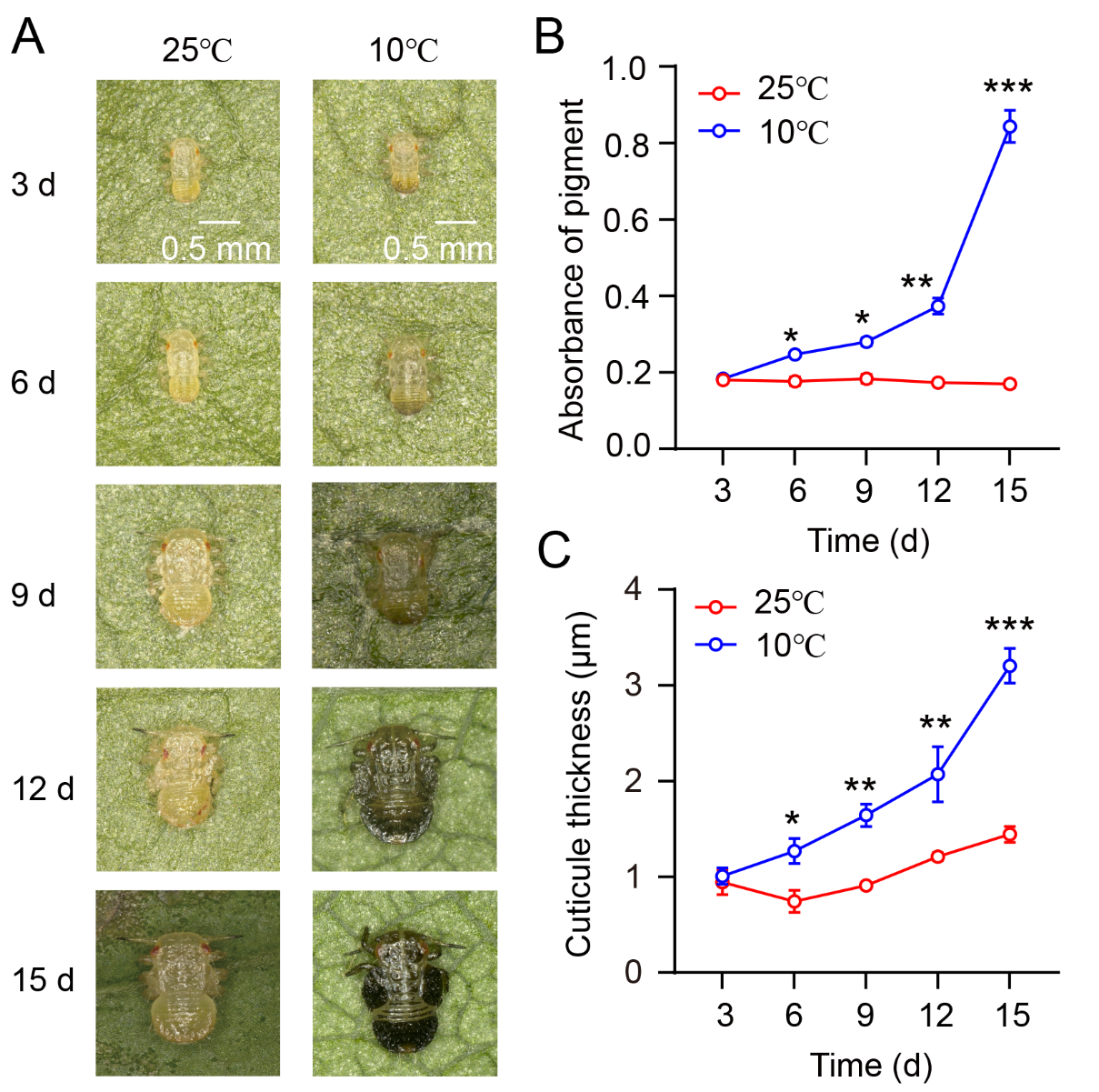

**Figure S1. Investigation of the relationship between nymph phenotype, cuticle pigment absorbance, and cuticle thickness during the transition from summer-form to winter-form in *C. chinensis*.**

(A) The nymph morphology during the transition from summer-form to winter-form of *C. chinensis* at varying time intervals (3, 6, 9, 12, 15 days). (B-C) The absorbance of total cuticle pigment and cuticle thickness of nymphs in *C. chinensis* under different temperature conditions.

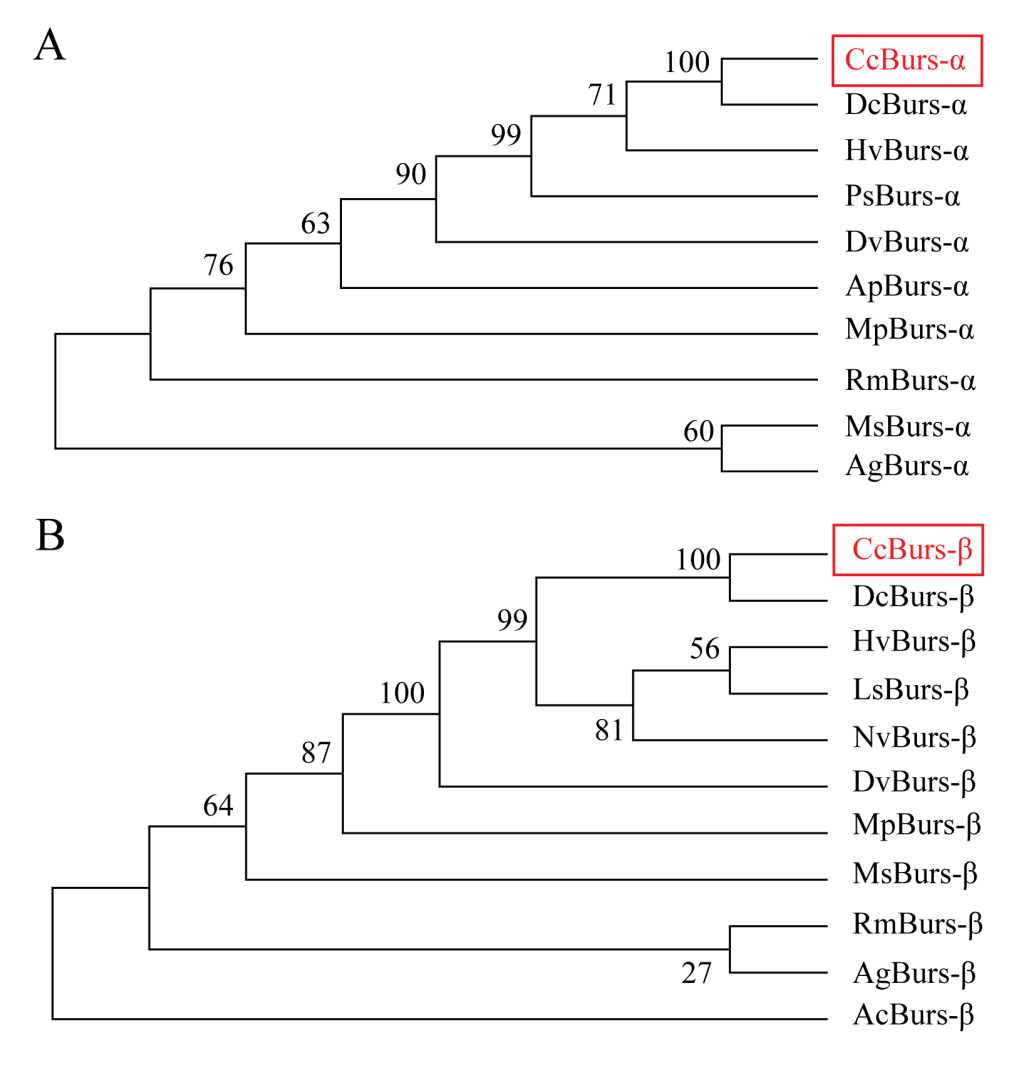

**Figure S2. Phylogenetic tree analysis of *CcBurs-α* and *CcBurs-β* with its homologs in other insect species.**

*HvBurs-α* (*Homalodisca vitripennis*, XP_046670477.1), *PsBurs-α* (*Phenacoccus solenopsis*, QEY08365.1), *DvBurs-α* (*Daktulosphaira vitifoliae*, XP_050533746.1), *RmBurs-α* (*Rhopalosiphum maidis*, XP_026815117.1), *MsBurs-α* (*Melanaphis sacchari*, XP_025201519.1), *AgBurs-α* (*Aphis gossypii*, XP_027848520.1). *DvBurs-β* (*D. vitifoliae*, XP_050533747.1), *MpBurs-β* (*Myzus persicae*, XP_022171709.1), *MsBurs-β* (*M. sacchari*, XP_025201526.1), *RmBurs-β* (*R. maidis*, XP_026815130.1), *AgBurs-β* (*A. gossypii*, XP_027848521.1), *AcBurs-β* (*Adelges cooleyi*, XP_050428069.1).

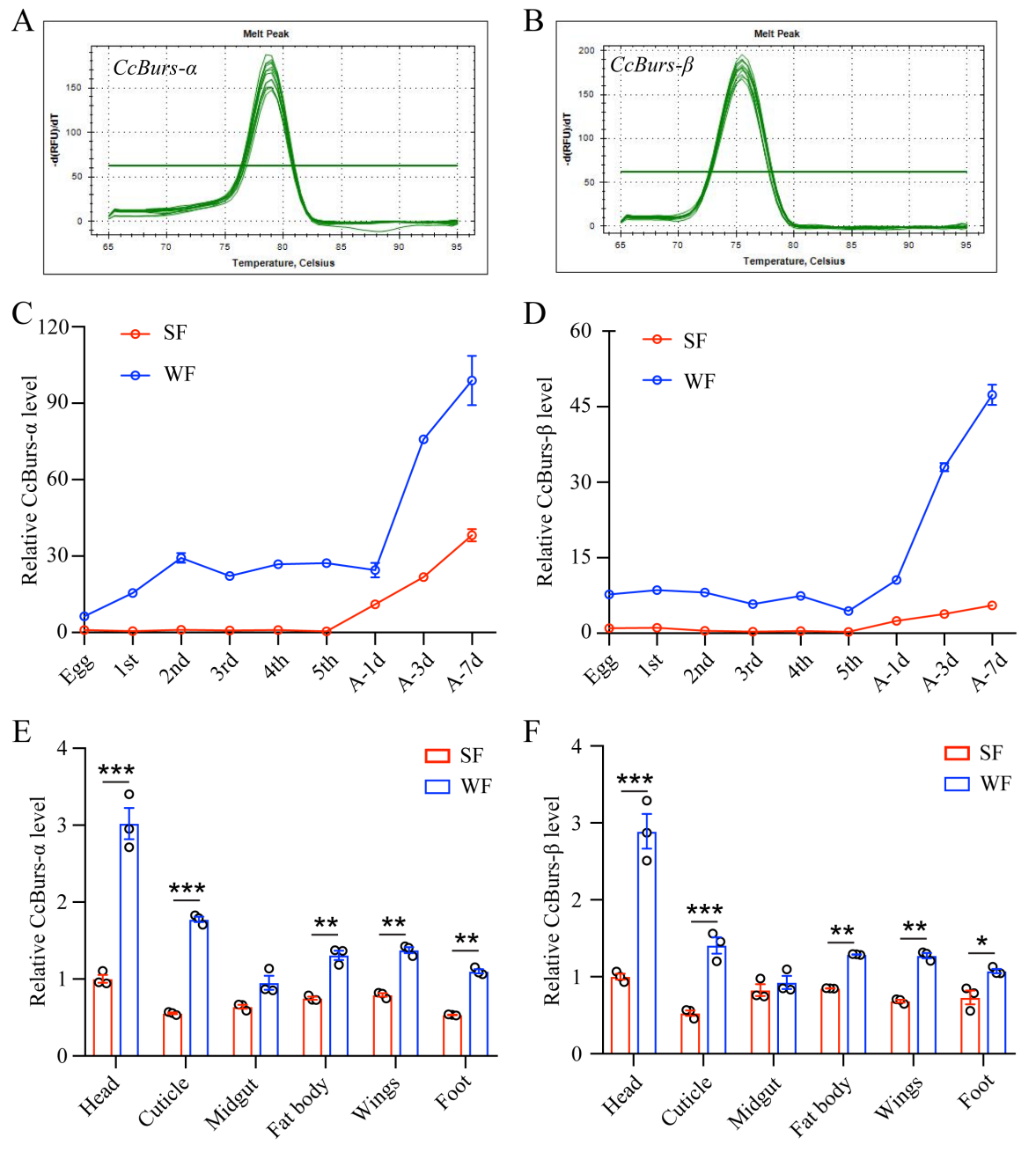

**Figure S3. Spatio-temporal expression patterns of *CcBurs-α* and *CcBurs-β.***

**A-B: Melting curve for qRT-PCR primers of *CcBurs-α* and *CcBurs-β*. C-D: The relative mRNA expression of *CcBurs-α* and *CcBurs-β* at different development ages of summer-form and winter-form by qRT-PCR (n=3).** 1st, 2nd, 3rd, 4th, and 5th are the nymphs at the first, second, third, fourth, and fifth instar, separately. A-1d, A-3d, and A-7d are the adults at the first day, third day, and seventh day, respectively.

**E-F: Tissue expression patterns of *CcBurs-α* and *CcBurs-β* in both summer-form and winter-form by qRT-PCR (n=3).** The data in S1C-S1F are shown as the mean ± SEM with three independent biological replications of at least 50 nymphs for each biological replication. Statistically significant differences were determined with the pair-wise Student's *t*-test in SPSS 20.0 software, and significance levels were denoted by * (*p* < 0.05), ** (*p* < 0.01), and *** (*p* < 0.001).

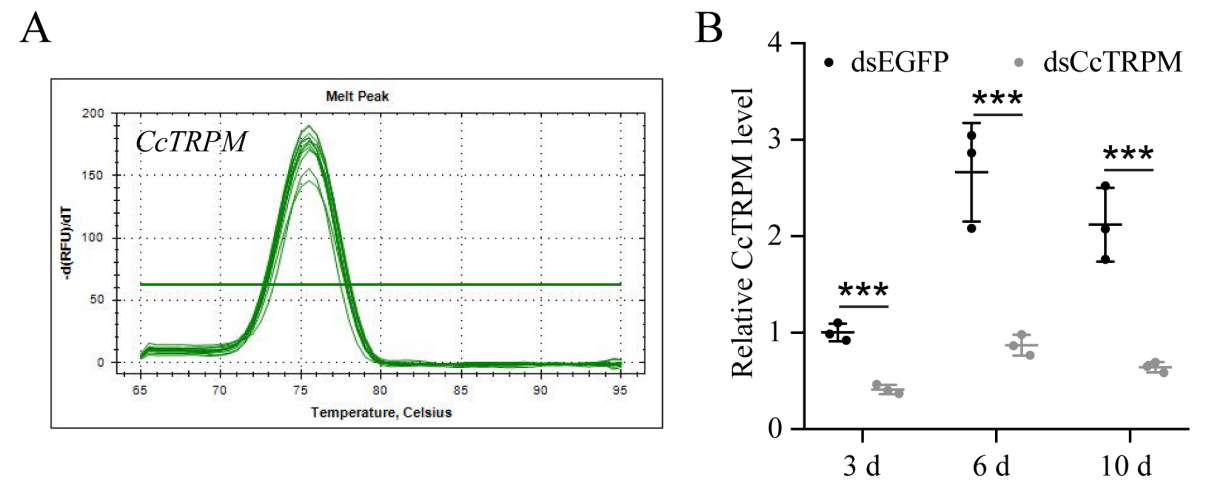

**Figure S4. RNAi efficiency of *CcTRPM* after dsRNA treatment at 3 d, 6 d, and 10 d by qRT-PCR under 10 °C condition (n=3).**

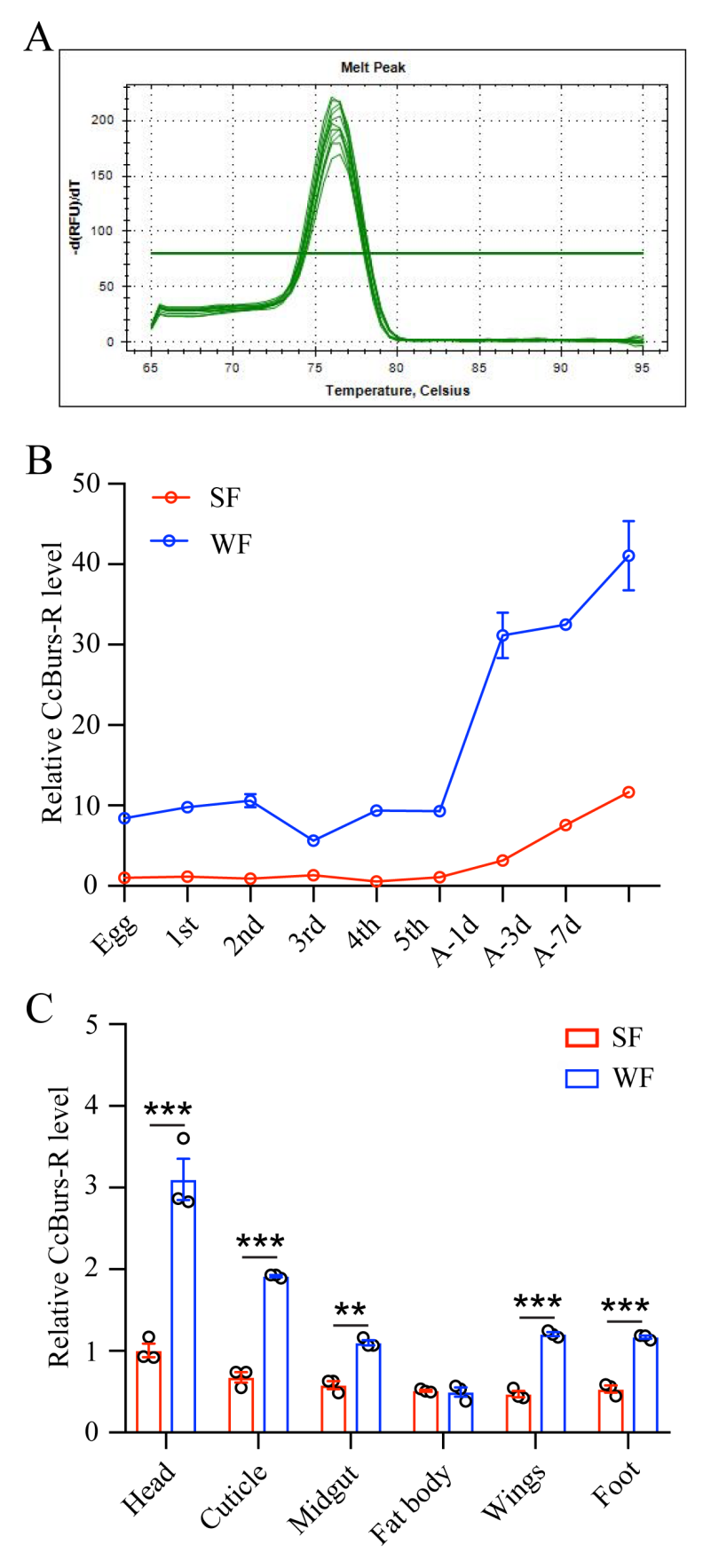

**Figure S5. Spatio-temporal expression patterns of *CcBurs-R* in both summer-form and winter-form by qRT-PCR (n=3).**

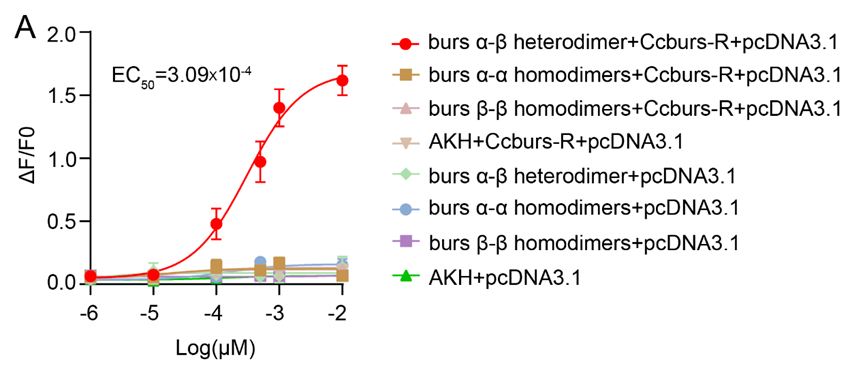

**Figure S6. Concentration-response relationships for peptides tested on *Ccburs-R*.**

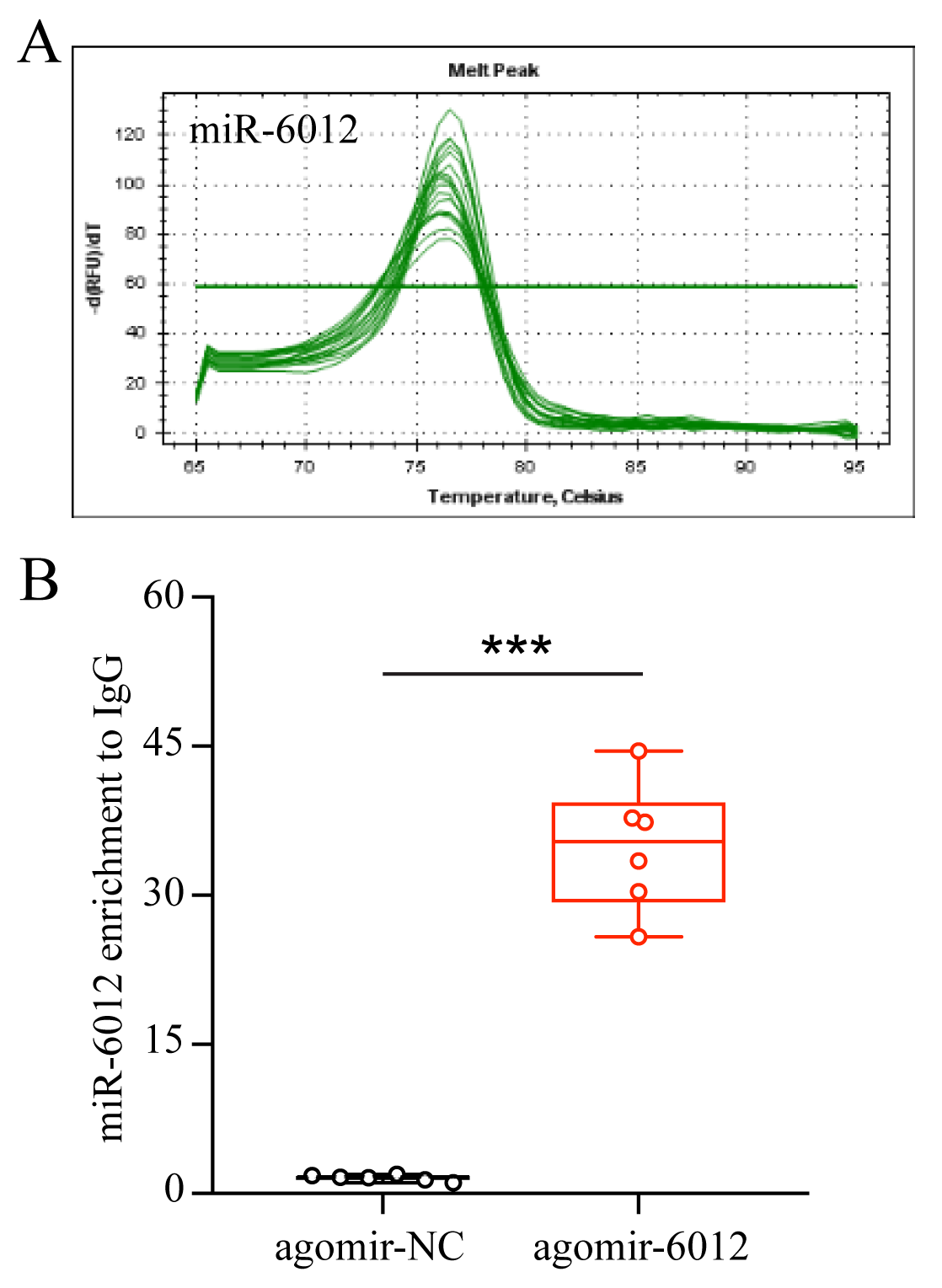

**Figure S7. Enrichment of miR-6012 by antibody against Ago1 in agomir-6012 treated group compared with agomir-NC group.**

**Table S1. List of primers used in this study.**

| **Gene name and GenBank ID** | **Forward Primer (5'-3')** | **Reverse Primer (5'-3')** | **Product size (bp)** | **Application purpose** |
| --- | --- | --- | --- | --- |
| *CcBurs-α*  (OR488624) | Full-F: CACATGTCGGACGGTATAA | Full-R: GCCACTCAATCTCTTCAGAC | 505 | Full-length cDNA cloning |
|  | RNAi-F1: ggatcctaatacgactcactatagg  AGTGTGTCTTCTGTTACCCCT | RNAi-R1: ggatcctaatacgactcactatagg  CACTAGCCTCTCTTTCTCCACT | 326 | dsRNA synthesis |
|  | qF: TACAAGCATTGAGGAGAG | qR: TCATTCTGACTTGGCAAA | 94 | qPCR |
|  | pcDNA3.1-his-mCherry-F1: ccactagtccagtgtggtggaattc  GGCCGCATCTTCTGGTAAA | pcDNA3.1-his-mCherry-R1: gtttaaacgggccctctagactcgag  CACTTAGTGATGGTGATGGTG | 114 | Reduced gel |
|  | pcDNA3.1-CcBurs-α-his-mCherry-F2: ctagcgtttaaacttaagctt  GCCACCATGTCGGACGGTATAATGAACA | pcDNA3.1-CcBurs-α-his-mCherry-R2: ccacactggactagtggatcc  TTCTGACTTGGCAAAATGTCC | 547 | Reduced gel |
|  | pcDNA3.1-CcBurs-α-his-P2A-mCherry-F3: gctggctagcgtttaaacttaagctt  GCCACCATGTCGGACGGTAT | pcDNA3.1-CcBurs-α-his-P2A-mCherry-R3: tccgcttccggtaccaagctt  GTGATGGTGATGGTGATGGTGGTG | 665 | Reduced gel |
|  | pcDNA3.1-CcBurs-α-his-P2A-mCherry-F1: TGGTGCACGTGGAAGCTTCCAACATTTCAGTCTCATCAGCC | pcDNA3.1-CcBurs-α-his-P2A-mCherry-R1: AGGCAGCTGGCGCGCTCCATGGTGGCAAGCTTAAGTTTAAAC | 6938 | Non-reduced gel |
|  | pcDNA3.1-CcBurs-α-his-P2A-mCherry-F2:  GCTGCTGCTGCTGCTCCCCCTGGTGCACGTGGAAGCTTCC | pcDNA3.1-CcBurs-α-his-P2A-mCherry-R2:  GGGGGAGCAGCAGCAGCAGCAGGCAGCTGGCGCGCTCCAT | 6958 | Non-reduced gel |
|  | Full-F: TTGTACACAAGCAATAATGTCC | Full-R: ACGTCATGAGTGACACAAAC | 446 | Full-length cDNA cloning |
| *CcBurs-β*  (OR488625) | RNAi-F1: ggatcctaatacgactcactatagg  TCCTTTTGTTTGAAATGGGCTCT | RNAi-R1: ggatcctaatacgactcactatagg  GGAAACTTTCTCGACAGCAGT | 336 | dsRNA synthesis |
|  | qF: GTGAGAAGATGGCAACATTAGA | qR: GCGAATAATCTCCACACTTGTA | 78 | qPCR |
|  | pcDNA3.1-P2A-mCherry-F1: tttaaacttaagcttggtacc  GGAAGCGGAGCTACTAACTTCAGCCTG | pcDNA3.1-P2A-mCherry-R1: gctcaccatggtggcggtacc  AGGTCCAGGGTTCTCCTCCACG | 108 | Reduced gel |
|  | pcDNA3.1-CcBurs-β-his-mCherry-F2: ctagcgtttaaacttaagctt  GCCACCATGTCCTTTTGTTTGAAATGGG | pcDNA3.1- CcBurs-β-his-mCherry-R2: ccacactggactagtggatcc  TCGCGAATAATCTCCACACT | 408 | Reduced gel |
|  | pcDNA3.1-CcBurs-β-his-P2A-mCherry-F3: gctggctagcgtttaaacttaagctt  GCCACCATGTCCTTTTGTTTGAAATGG | pcDNA3.1-CcBurs-β-his-P2A-mCherry-R3: tccgcttccggtaccaagctt  GTGATGGTGATGGTGATGGTGGTG | 590 | Reduced gel |
|  | pcDNA3.1-CcBurs-β-his-P2A-mCherry-F1: TGGTGCACGTGGAAGCTTCCGAAAAGGACGAGGCTTGT | pcDNA3.1-CcBurs-β-his-P2A-mCherry-R1:  AGGCAGCTGGCGCGCTCCATGGTGGCAAGCTTAAGTTTAAAC | 6863 | Non-reduced gel |
|  | pcDNA3.1-CcBurs-β-his-P2A-mCherry-F2:  GCTGCTGCTGCTGCTCCCCCTGGTGCACGTGGAAGCTTCC | pcDNA3.1-CcBurs-β-his-P2A-mCherry-R2:  GGGGGAGCAGCAGCAGCAGCAGGCAGCTGGCGCGCTCCAT | 6883 | Non-reduced gel |
| *CcBurs-R*  (OR488626) | Full-F: TTTCATCGTCGTTTGGAGAT | Full-R: CCCGAAAGTATCCCATAGTG | 1758 | Full-length cDNA cloning |
|  | RNAi-F1: ggatcctaatacgactcactatagg  ATCTGACCCAATTGAGGAAC | RNAi-R1: ggatcctaatacgactcactatagg  CCGTCCGGTATGTATTTGAT | 371 | dsRNA synthesis |
|  | orf-F1 | orf-R1 |  |  |
|  | AGGTCTATATAAGCAGAGCTC | AGGTCTATATAAGCAGAGCTC | 1811 | Amplification of |
|  | ATGAAGAAAACGTCATTTTTAATA | ATCAATACCCGGAGTCAAATGA |  | orf sequence |
|  | qF: CCTCAGCAAACTACAAGTCT | qR: GCCCAGGTTCAAATCTTCTA | 103 | qPCR |
|  | 3'UTR-Full-F1: tctagttgtttaaacgagctc  CGCTTGGCATAGTTCCATAT | 3'UTR-Full-R1: cctgcaggtcgactctaga  GGTACCTAACCTACCCAGAT | 599 | Application of  3'UTR Full sequence |
|  | 3'UTR-Mut-F1: tctagttgtttaaacgagctc  TTGTTCCCACTTCTTCCTGGAA | 3'UTR-Mut-R1: cctgcaggtcgactctaga  GGTACCTAACCTACCCAGAT | 382 | Application of  3'UTR mutant sequence |
| *CcTRPM*  (OQ658558) | RNAi-F: ggatcctaatacgactcactatagg  GGTACCTTCGTATCCTCAAC | RNAi-F: ggatcctaatacgactcactatagg  TGGCATTGATCTCCATGAAA | 472 | dsRNA synthesis |
|  | qF: ACAATGCATTCTTCTTGACC | qR: GACTTTGTACACTGGGGGTA | 141 | qPCR |
| *EGFP*  (ACY56286) | RNAi-F: ggatcctaatacgactcactatagg  ACTCCAGCAGGACCATGTGATC | RNAi-F: ggatcctaatacgactcactatagg  CCTGAAGTTCATCTGCACCAC | 596 | dsRNA synthesis |
| *CcTre1*  (OQ734934) | qF: GGAACTCCCTCCTCTATGTT | qR: CCAATAGTCAGCCAATCTGT | 147 | qPCR |
| *CcCHS1*  (OQ658570) | qF: AGAAGAGAAGAAACAGCAGG | qR: TAGTGTCCAATCGTTTCTCC | 233 | qPCR |
| *Ccβ-actin*  (OQ658571) | qF: CGTATGCAGAAGGAAATCAC | qR: AGATCCACATCTGTTGGAAG | 139 | qPCR |
| miR-6012 | qF: TTCGGCGATGAGATCAGCCAGT | - | - | qPCR |
| U6 | qF:AGGATGACACGCAAAATCGT | - | - | qPCR |

**Table S2. Comparison of pigmentation and cuticule thickness after *CcTRPM*, *CcBurs-a*, *CcBurs-β*, and *CcBurs-R* knockdown.**

|  | **dsEGFP** | **dsCcTRPM** | **dsCcBurs-a** | **dsCcBurs-β** | **dsBurs-R** |
| --- | --- | --- | --- | --- | --- |
| Pigmentation  (absorbance at 300nm) | 0.8499 ± 0.0532a | 0.1779 ± 0.0059b | 0.1846 ± 0.0203b | 0.1931 ± 0.0309b | 0.1410 ± 0.0131b |
| Cuticule thickness  (μm) | 3.39 ± 0.20a | 1.75 ± 0.09b | 1.44 ± 0.12b | 1.54 ± 0.15b | 1.34 ± 0.13b |
